# PI5P4Kα Regulates Iron Balance to Promote Metabolic Fitness in Pancreatic Cancer

**DOI:** 10.1101/2025.04.07.647654

**Authors:** Gurpreet K. Arora, Ryan M. Loughran, Kyanh Ly, Cheska Marie Galapate, Alicia Llorente, Taylor R. Anderson, Chantal Pauli, David A. Scott, Yoav Altman, Rabi Murad, Cosimo Commisso, Brooke M. Emerling

## Abstract

Phosphoinositide kinases generate distinct phosphoinositides that regulate processes that maintain cellular fitness. Phosphatidylinositol 5-phosphate 4-kinases (PI5P4Ks) have garnered interest for their role in cancer metabolism and cellular trafficking; however, their function in pancreatic ductal adenocarcinoma (PDAC) remains unexplored. Given the unique metabolic demands of PDAC cells, which heavily rely on altered trafficking pathways to support their growth, investigating PI5P4Ks in this context may reveal critical insights. We identify PI5P4Kα as a regulator of PDAC cell fitness through its key role in maintaining iron homeostasis. PI5P4Kα depletion causes metabolic disruptions and reduced intracellular iron import, leading to the induction of apoptosis in PDAC cells that is reversed by iron supplementation. Notably, we find that PI5P4Kα knockdown suppresses tumor growth in a xenograft mouse model of PDAC. These results not only illuminate the mechanistic underpinnings of PI5P4Kα function in PDAC but also position it as a promising therapeutic target for this disease.

**Graphical Abstract:** 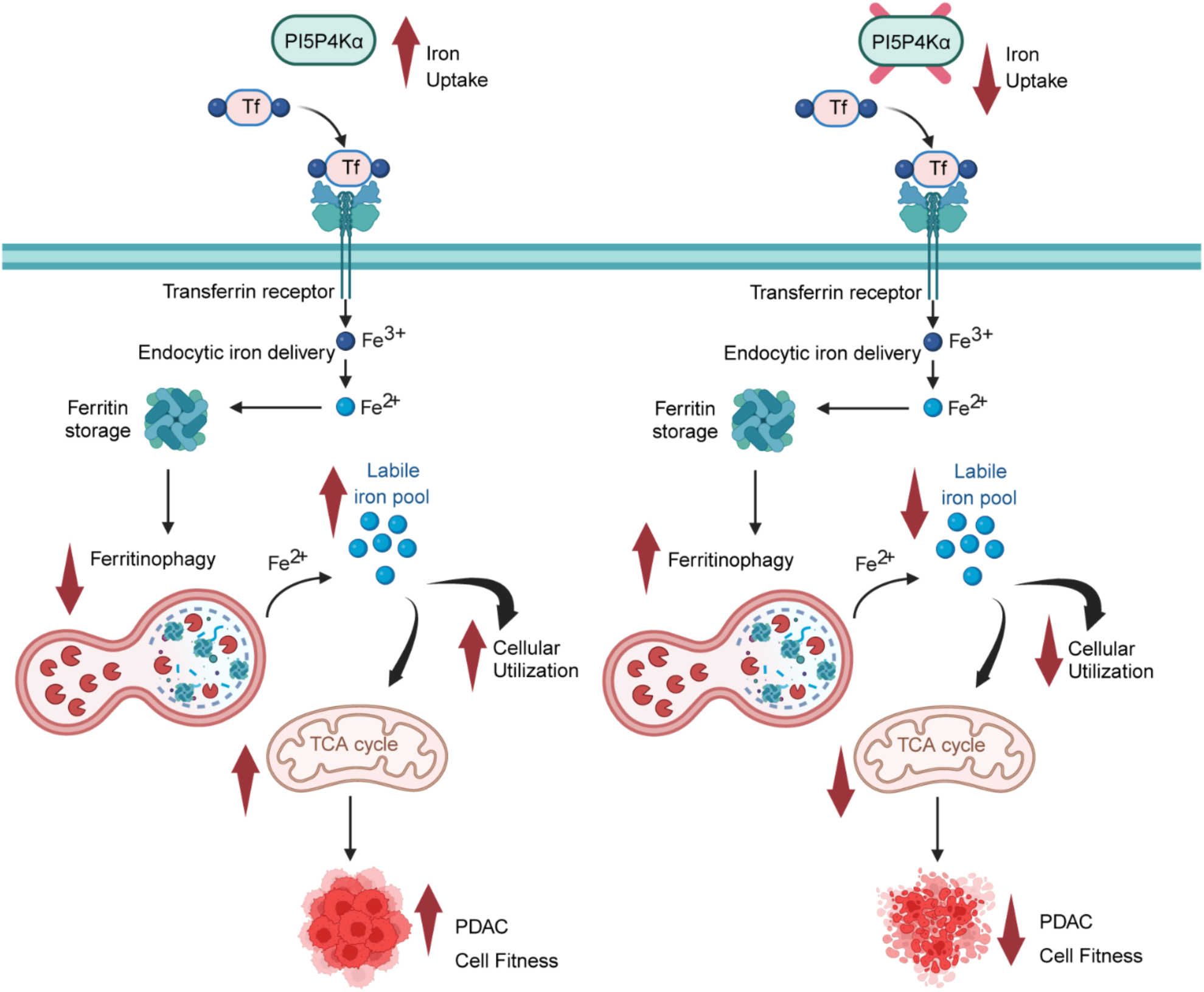

**Schematic representation of the novel function and implication of PI5P4Kα in PDAC**. PI5P4Kα is upregulated in PDAC to support the ferro-addiction and heightened metabolic requirements of the tumor. When PI5P4Kα is inhibited, the PDAC cells undergo apoptotic cell death due to iron depletion. PI5P4Kα-depleted PDAC cells upregulate autophagy as an adaptive response but is not sufficient to defend against apoptotic cell death.

## Introduction

Phosphoinositide kinases are a specialized family of lipid kinases that phosphorylate inositol lipids and their derivatives to generate seven unique phosphoinositide species with spatial and temporal precision^1,2^. Phosphoinositides are critical regulators of a plethora of cellular processes including membrane trafficking, cellular signaling and metabolism, many of which are frequently dysregulated to support tumorigenesis^3–5^. Among the phosphoinositide kinases, phosphoinositide 3-kinases (PI3Ks) have long been known to drive multiple cancer types due to gain-of-function mutations leading to the activation of AKT signaling^6^. PI3Ks have consequently been the most extensively studied and clinically investigated lipid kinases as evidenced by a myriad of clinical trials. However, toxicity concerns and resistance mechanisms have dampened the clinical success of inhibitors targeting PI3K and its downstream signaling^7^.

Recently, there has been a shift in efforts toward the less studied phosphoinositide kinases^1^, including the phosphatidylinositol phosphate (PIP) kinase family member phosphatidylinositol 5-phosphate 4-kinase (PI5P4K), which primarily phosphorylates the lipid phosphatidylinositol 5-phosphate, PI(5)P, to generate phosphatidylinositol 4,5-bisphosphate PI(4,5)P2^8–10^.PI5P4Ks have been implicated in various cancers and our recent studies have identified important roles for these enzymes in cellular metabolic processes such as stress responses^11–13^, cellular trafficking^14^ and organelle communication^15^. The PI5P4K family consists of three isoforms, PI5P4Kα, PI5P4Kβ, and PI5P4Kγ, with PI5P4Kα being the most catalytically active isoform. The knockout mouse model for the PI5P4Kα isoform has been generated and is viable with normal life span^11^, emphasizing that targeting PI5P4Kα in cancer may not be deleterious to normal cells, which is a concern with some of the other phosphoinositide kinases such as the PI4K family^1^. Interestingly, the *PIP4K2A* gene (encoding PI5P4Kα) is often transcriptionally upregulated in tumor samples from patients in multiple cancer types, underscoring its role in supporting the elevated metabolic demands of tumor cells^16^. Importantly, targeting PI5P4Kα has been shown to be a mutant p53-dependent susceptibility in breast cancer^11,16,17^. Given the increasing links between PI5P4Ks and cancer progression, there has been a recent translational interest in the early development of inhibitors targeting specific PI5P4Ks^1,18^.

Pancreatic ductal adenocarcinoma (PDAC) is characterized by unique metabolic dependencies that support its aggressive growth, often relying on altered or enhanced trafficking pathways such as macropinocytosis and autophagy that fuel tumor survival^19–23^. Furthermore, the majority of PDAC tumors harbor mutations in the tumor suppressor gene p53, which exacerbates these metabolic alterations^24,25^. Whether PI5P4Ks represent a therapeutic vulnerability in PDAC has not been investigated. Here, we report the discovery that PI5P4Kα is an essential metabolic dependency in PDAC cells and intriguingly this dependency is due to the novel role of PI5P4Kα in iron homeostasis. Depleting PI5P4Kα induces apoptotic cell death selectively in PDAC cells, particularly those with *TP53* mutations. This vulnerability arises from a disruption in intracellular iron homeostasis and a diminishment in TCA cycle intermediates, whose production by TCA cycle enzymes is known to be dependent on intracellular iron^26,27^. Consistent with these observations, we find that exogenous iron rescues the deleterious effects of PI5P4Kα suppression on cell fitness. PDAC cells have been shown to maintain iron balance via ferritinophagy, a form of selective autophagy used to release iron^28,29^ phosphatidylinositol 3-phosphate; however, we find that instead PI5P4Kα controls iron homeostasis through transferrin-bound iron import. Emphasizing the physiological relevance of PI5P4Kα in PDAC, we find that *PIP4K2A* expression is upregulated in human PDAC specimens and correlates with multiple iron uptake gene

signatures. High expression of *PIP4K2A*, as well as of genes in the signatures for iron import and transport, are associated with poor prognosis in PDAC patients. Moreover, using a xenograft mouse model of PDAC, we determine that PI5P4Kα knockdown significantly suppresses tumor growth. Altogether, our work exposes a novel homeostatic function of PI5P4Kα as a mediator of iron balance, a role that has not been shown before for any other PIP kinase member^8^, and reveals that the targeting of PI5P4Ks in PDAC may represent a novel therapeutic modality for this disease.

## Results

### PI5P4Kα depletion triggers apoptotic cell death selectively in PDAC cells

To evaluate the effects of PI5P4Kα suppression on the fitness of pancreatic ductal adenocarcinoma (PDAC) cells, we utilized shRNA-mediated knockdown to deplete PI5P4Kα in MIA PaCa-2 and PANC-1 cells. To evaluate whether any observed effects could be selective to cancer cells, we also knocked down PI5P4Kα in HPNE cells, an untransformed epithelial cell line derived from normal human pancreas. Cells were infected with either a non-targeting control shRNA (sh scr) or with two distinct shRNAs targeting *PIP4K2A* (sh α#1 and sh α#2). To assess knockdown efficiency, we performed qRT-PCR, which demonstrated a significant decrease in *PIP4K2A* transcripts in both PDAC cell lines relative to sh scr control (Figure S1A). To investigate effects on cell fitness, relative cell number was evaluated following knockdown using a crystal violet staining assay. Knockdown of PI5P4Kα led to a marked reduction in cell number in both MIA PaCa-2 and PANC-1 cells (Figures 1A and 1B). In addition, a time-course experiment revealed a cytotoxic effect on the PDAC cells upon PI5P4Kα depletion (Figure S1B). Notably, this cytotoxic effect was selective to PDAC cells, as depletion of PI5P4Kα in HPNE cells did not impact cell number (Figures 1A and 1B and Figure S1B).

**Figure 1.**
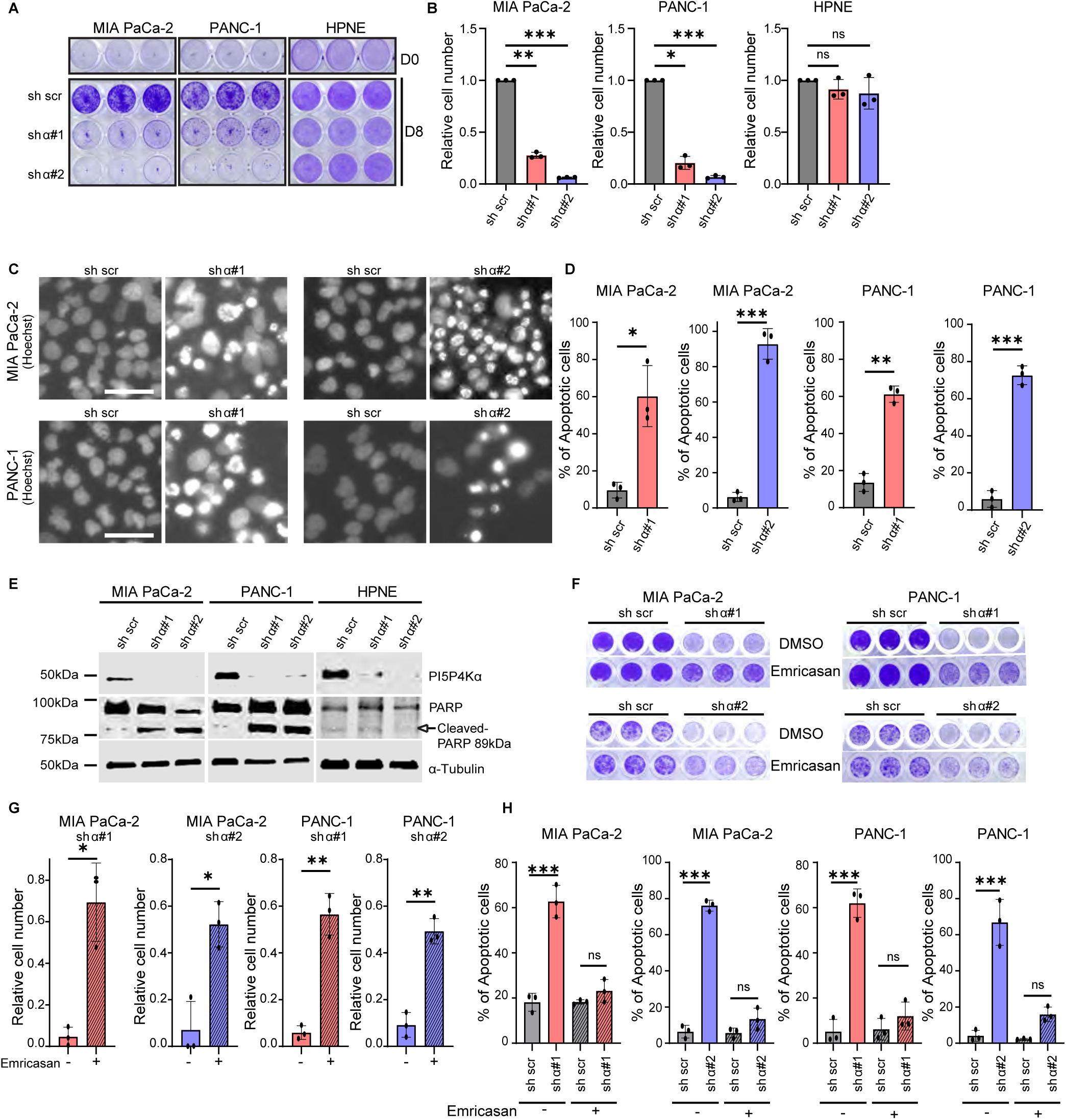
Knockdown of PI5P4Kα induces apoptotic cell death in PDAC cells. **(A-B)** Crystal violet staining and quantification of relative cell number in MIA PaCa-1, PANC-1 or HPNE cells, trans-duced with nontargeting negative control sh scr or two different hairpins targeting PIP4K2A (sh α#1 or sh α#2). **(A)** Representative images are shown for Day 0 (D0) and Day 8 (D8) post-transduction. **(B)** Quantification of relative cell number using the crystal violet stained area in the indicated cell lines D8 post-transduction. Data is presented relative to the sh scr condition and is the average of three independent experiments. **(C-D)** Hoechst staining assay in the indicated cell lines transduced with sh scr, sh α#1 or sh α#2. **(C)** Representative images are shown. (scale bar, 50µM). **(D)** Quantification of the percentage of apoptotic cells. Data is the average of three independent experiments. **(E)** Immunoblot assessing PI5P4Kα and cleaved PARP protein levels in the indicated cell lines transduced with sh scr, sh α#1 or sh α#2. Cells were harvested 3 days post-transduction for sh α#2 and 6 days post-transduction for sh α#1. Tubulin was used as a loading control. Data is representative of three independent experiments. **(F-H)** Crystal violet staining, quantification of relative cell number and Hoechst staining in the indicated cell lines transduced with sh scr, sh α#1 or sh α#2 in the absence or presence of Emricasan. **(F)** Representative image of crystal violet assays. **(G)** Quan-tification of crystal violet stained area. Data is presented relative to the sh scr condition and is the average of three independent experiments. **(H)** Quantification of the percentage of apoptotic cells. Data is the average of three inde-pendent experiments. **(B, D, G, H)** Data is presented as the mean ± sd. Statistical significance was calculated using unpaired two-tailed Student’s t-test. h, Statistical significance was calculated using one-way Anova. ns – not significant, *P < 0.05, **P ≤ 0.001, ***P ≤ 0.0001.

To investigate whether the observed loss of PDAC cell fitness caused by PI5P4Kα depletion was a result of apoptosis, we utilized a Hoechst nuclear staining assay since apoptotic cells display nuclear condensation and DNA fragmentation^30^. Our analysis revealed a significant increase in the percentage of apoptotic cells following depletion of PI5P4Kα (Figures 1C and 1D). To further validate that PDAC cells were indeed undergoing apoptosis, we performed western blot analysis to evaluate the levels of cleaved poly (ADP-ribose) polymerase (PARP), as its cleavage by caspases serves as a hallmark of the final stages of apoptosis. In accordance with our previous results, PI5P4Kα-depleted cells displayed enhanced PARP cleavage, resulting in the increased levels of the 89 kDa cleavage fragment (Figure 1E). Importantly, knockdown of PI5P4Kα in the normal HPNE cells did not result in increased PARP cleavage (Figure1E). Directly linking the observed cytotoxic effects of PI5P4Kα depletion to apoptosis, we found that treatment with Emricasan, a pan-caspase inhibitor^31^, rescued both cell fitness (Figures 1F and 1G), the extent of apoptosis in PI5P4Kα-depleted cells (Figure1H and Figure S1C), and prevented the cleavage of PARP (Figure S1D).

MIA PaCa-2 and PANC-1 cells both exhibit mutations in the tumor suppressor gene *TP53*, which are observed in more than 60% of PDAC patients^32^. We previously reported that PI5P4K is particularly important for tumor cell viability in breast cancer cells harboring *TP53* mutations^11^. To test whether this might also be the case in PDAC, we knocked down PI5P4Kα in SW1990 and HPAF-II cells, which lack inactivating mutations in *TP53*. We found that PI5P4Kα depletion had no effect or led to a moderate decrease in proliferation in these cells, and we did not observe any evidence of apoptosis (Figures S1E and S1F). Altogether, these findings suggest a crucial role for PI5P4Kα in controlling cell fitness in PDAC cells, and the differential response observed with varying *TP53* status highlights a conditional essentiality of PI5P4Kα in PDAC.

### PI5P4Kα functions to maintain intracellular levels of TCA cycle intermediates

Given that PI5P4Kα is contextually crucial for cellular energetics and that apoptosis can be induced by metabolic stress, we next sought to assess the impact of PI5P4Kα suppression on the cellular metabolism of PDAC cells. We performed targeted metabolomics utilizing gas chromatography-mass spectrometry (GC/MS). We measured and quantified intracellular polar metabolites in PI5P4Kα-knockdown cells relative to control cells. The most significantly decreased metabolites in PI5P4Kα-depleted cells were α-ketoglutarate (α-KG), fumarate, and malate, all of which are key intermediates of the tricarboxylic acid (TCA) cycle (Figure 2A) We also observed a significant decrease in lactate, and to a lesser extent, pyruvate (Figure 2B). To gain deeper insights into these metabolic effects, we conducted heavy isotope tracing studies using either ^13^C5-glutamine or ^13^C6-glucose as substrates (Figure 2C). Interestingly, we found a shift in the labeling of TCA cycle metabolites in PI5P4Kα-depleted cells, with less labeling with ^13^C5-glutamine and proportionally more labeling with ^13^C6-glucose (Figures 2D and 2E). These data suggested that the alterations in the TCA cycle or glycolytic metabolites could be responsible for the cell fitness defects observed upon PI5P4Kα knockdown in PDAC cells.

**Figure 2.**
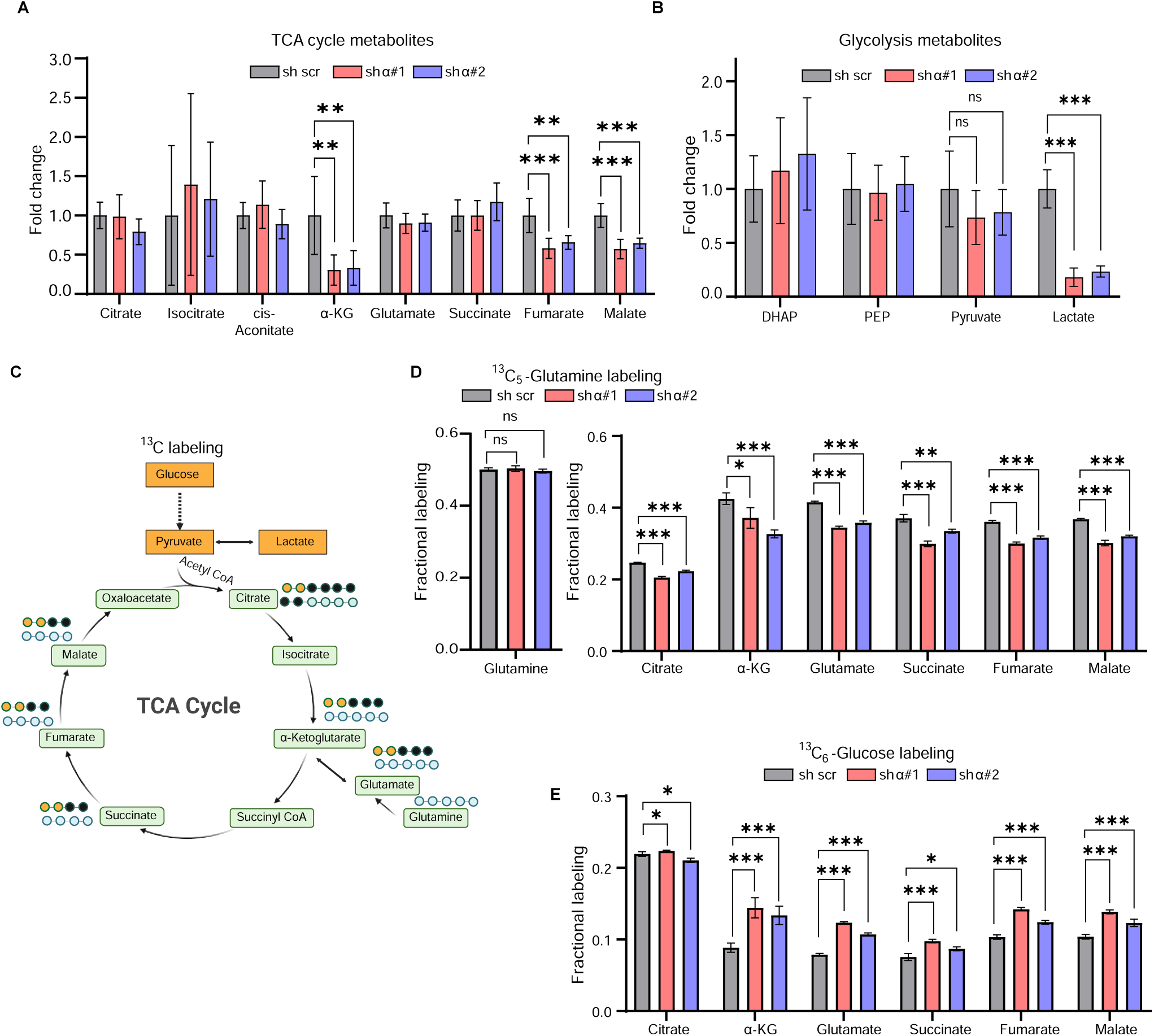
PI5P4Kα inhibition triggers metabolic perturbations. **(A-B)** Quantification of intracellular levels of the metabolites in MIA PaCa-2 cells transduced with sh scr, sh α#1 or sh α#2. Cells were harvested 3 days post-transduction. Data is presented relative to the sh scr condition and is a repre-sentative of two independent experiments. Data are presented as mean ± sd of n = 10 samples. **(A)** Quantification of TCA cycle metabolites. **(B)** Quantification of glycolysis metabolites. **(C-E)** Labeling of TCA cycle from 13C6-glucose or 13C5-glutamine. **(C)** Schematic representation of 13C labeling. 13C5-glutamine will predominantly label four or five carbons in the TCA cycle, as indicated by the light blue circles. 13C6-glucose, metabolized via acetyl-CoA, will predominantly label two carbons in the TCA cycle, as indicated by the orange circles. (Unlabeled carbons shown in black). **(D)** Fractional labeling from 13C5-glutamine. **(E)** Fractional labeling from 13C6-glucose. Data are presented as mean ± sd of n = 5 samples **(D-E)**. Statistical significance was calculated using unpaired two-tailed Student’s t-test. ns – not significant, *P < 0.05, **P ≤ 0.001, ***P ≤ 0.0001.

To ascertain which metabolic perturbations were mediating the apoptotic response upon PI5P4Kα knockdown, we performed orthogonal rescue assays using different metabolites. We found that exogenous supplementation with either pyruvate or lactate failed to protect the PI5P4Kα-depleted PDAC cells from undergoing apoptosis (Figure 3A). To investigate whether the observed cell death resulting from PI5P4Kα knockdown is linked to the reductions in the TCA cycle metabolites, we conducted rescue assays with dimethyl α-ketoglutarate (DMKG) or dimethyl DL-glutamate (DMGlu), cell permeable versions of α-KG and glutamate, respectively. Glutamate fuels the TCA cycle since in the early steps of glutaminolysis it is converted to α-KG, a critical TCA cycle intermediate. Interestingly, supplementation with these exogenous metabolites rescued cell fitness in PI5P4Kα-deficient cells (Fig. 3b), suppressed apoptosis (Figures 3C-3E) and reduced PARP cleavage (Figures 3F and 3G). Taken together, these findings indicate that PI5P4Kα functions to support central carbon metabolism by maintaining the intracellular levels of the TCA cycle metabolites.

**Figure 3.**
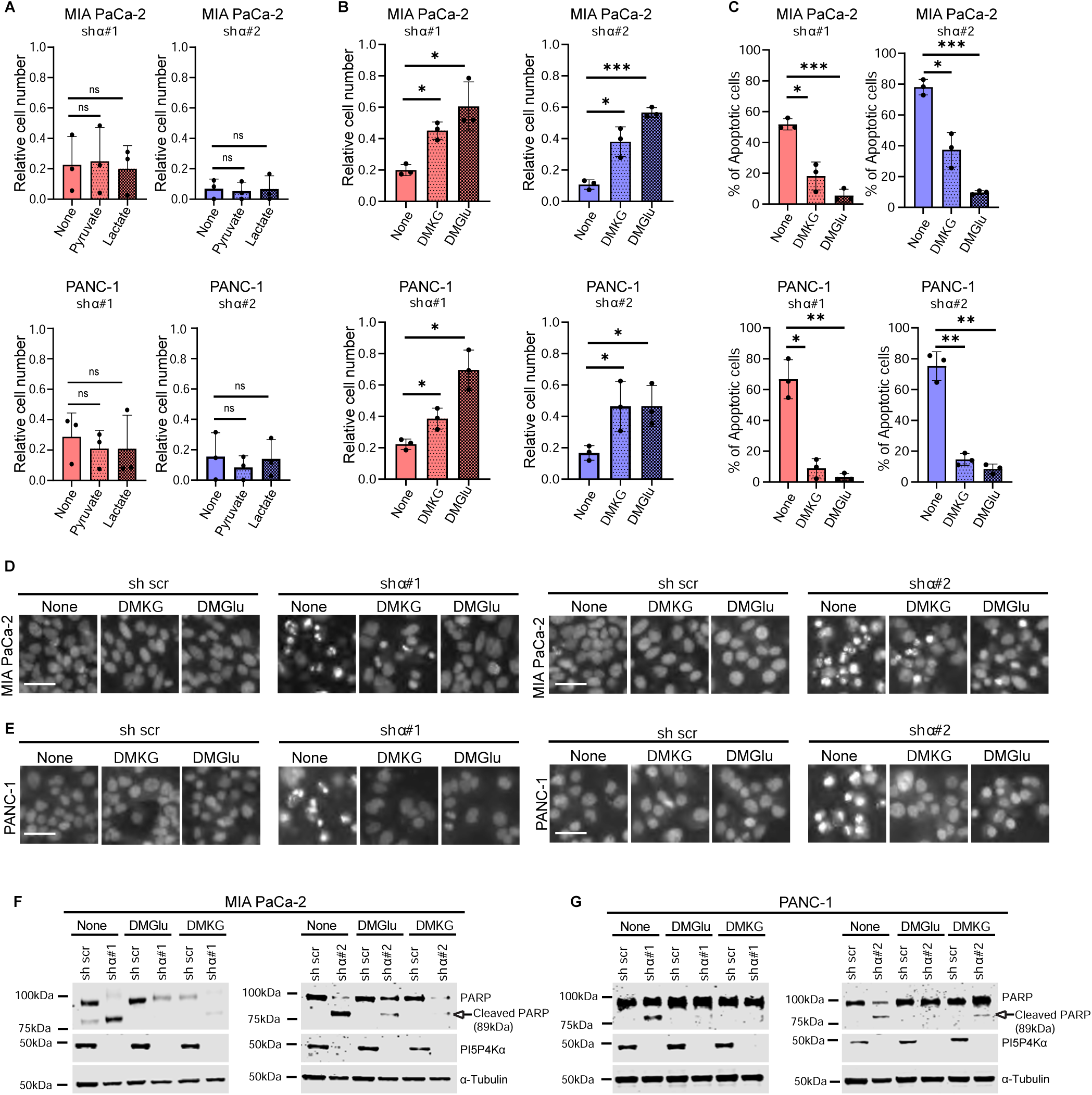
TCA metabolites rescue cell fitness effects of PI5P4Kα-depletion in PDAC cells. **(A-B)** Quantification of relative cell number using the crystal violet stained area in the indicated cell lines transduced with sh scr, sh α#1 or sh α#2 in the absence or presence of pyruvate, lactate **(A)** Dimethyl α-ketoglutarate (DMKG) or Dimethyl DL-Glutamate hydrochloride (DMGlu) **(B)**. Data is presented relative to the sh scr condition for each respec-tive treatment and is the average of three independent experiments. **(C-G)** Hoechst staining assay and immunoblots in the indicated cell lines transduced with sh scr, sh α#1 or sh α#2 in the absence or presence of DMKG or DMGlu. **(C)** Quantification of the percentage of apoptotic cells. Data is the average of three independent experiments. **(D)** Representative images of Hoechst staining in MIA PaCa-2. **(E)** Representative images of Hoechst staining in PANC-1 (scale bar, 50µM). Immunoblot assessing PI5P4Kα and cleaved PARP protein levels in MIA PaCa-2 **(F)** and PANC-1 **(G)**. Tubulin was used as a loading control. Data is representative of three independent experiments. **(A, B, C)** Data are presented as the mean ± sd. Statistical significance was calculated using unpaired two-tailed Student’s t-test. ns – not significant, *P < 0.05, **P ≤ 0.001, ***P ≤ 0.0001.

### PI5P4Kα regulates PDAC cell fitness through iron homeostasis

We next sought to decipher how PI5P4Kα is impacting TCA cycle metabolite levels. Many enzymes that are integral to the TCA cycle utilize iron-containing heme as a prosthetic group or rely on iron-sulfur clusters as essential cofactors^27,28,33^. In addition, recent studies have revealed the relevance of iron metabolism in PDAC, where both deficiency and excess of the intracellular iron pools can affect PDAC cell viability^26,28,34,35^. Considering these connections and our observed metabolic changes, we investigated intracellular iron levels in PI5P4Kα-knockdown cells. Initially, we investigated intracellular iron levels using the probe FerroOrange, which selectively reacts with Fe^2+^ but not Fe^3+^. We observed reduced FerroOrange signal intensity in PI5P4Kα-knockdown cells in the PDAC cells, indicative of cytoplasmic iron imbalance (Figures 4A and 4B). To determine whether this iron imbalance plays a role in PI5P4Kα-mediated cell fitness, we supplemented the growth medium with iron in the form of ferric (III) ammonium citrate (FAC), which restores intracellular iron (Figure4B, Figure S2A). Remarkably, FAC supplementation rescued the deleterious effects of PI5P4Kα depletion on cell fitness, reversing both cell growth defects and apoptosis-driven production of cleaved PARP (Figures 4C-4G). To rule out the possibility that the rescue effects are attributed to the citrate component of FAC, we supplemented the growth medium with sodium citrate, which did not rescue apoptosis (Figures S2B and S2C).

**Figure 4.**
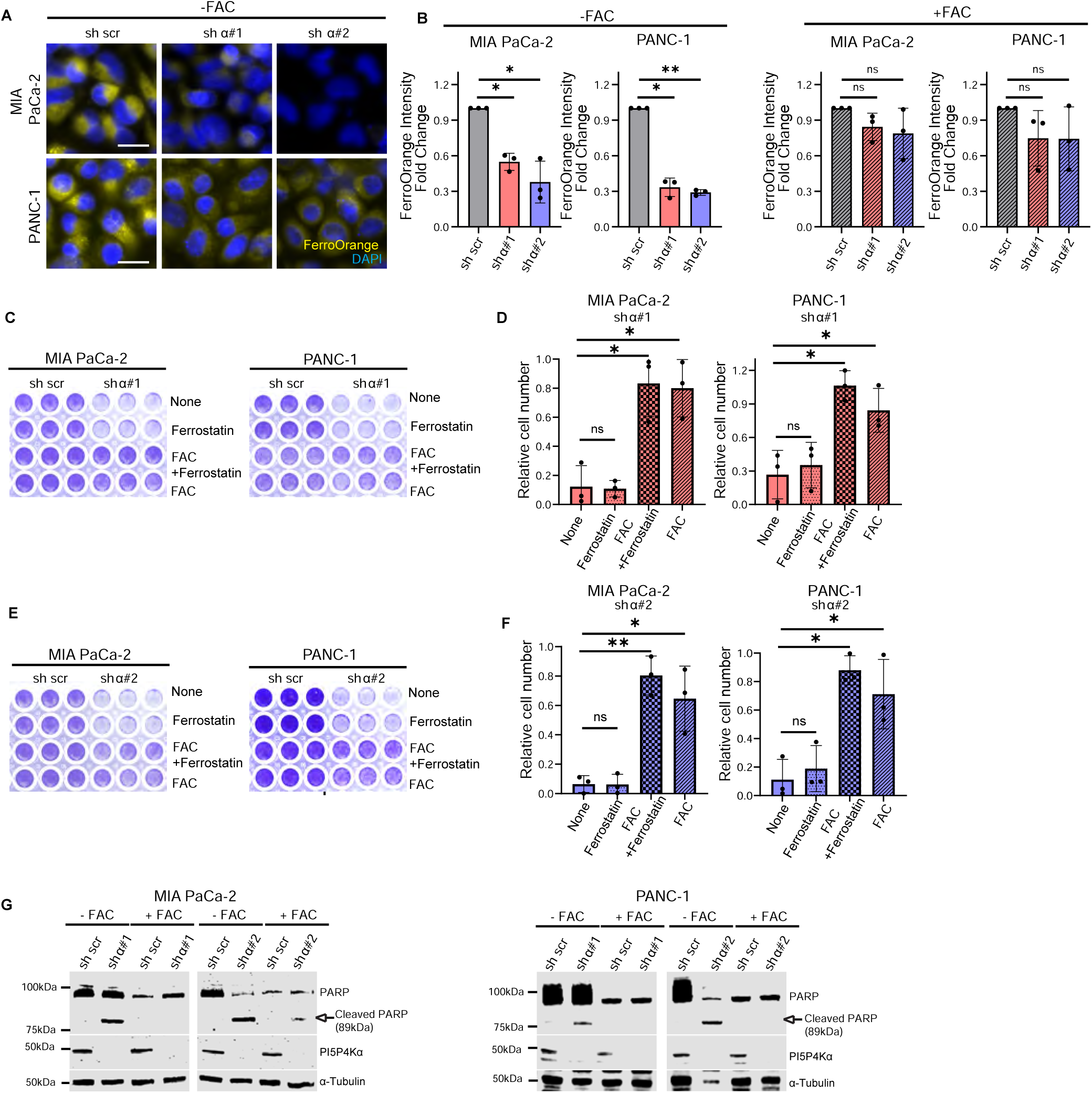
PI5P4Kα maintains intracellular levels of iron to promote PDAC cell fitness. **(A-B)** FerroOrange staining in the indicated cell lines transduced with sh scr, sh α#1 or sh α#2 without FAC **(A-B)** and with FAC **(B)**. Cells were imaged between 3-4 days post-transduction. **(A)** Immunofluorescence microscopy images of FerroOrange and DAPI indicating the representative effect of three independent experiments (scale bar, 20 µm). **(B)** Quantitation of immunofluorescence microscopy images. Data is presented as signal intensity relative to the sh scr condition and is the average of three independent experiments. **(C-F)** Crystal violet staining and quantification of relative cell number in the indicated cell lines transduced with sh scr, sh α#1 or sh α#2 in the absence or presence of Ferrostatin, Ferrostatin with ferric ammonium citrate (FAC) and FAC alone. Data is presented relative to the sh scr condition for each respective treatment and is the average of three independent experiments. **(C)** Representative image of crystal violet assay of cell lines transduced with sh scr or sh α#1. **(D)** Quantification of crystal violet stained area. **(E)** Representative image of crystal violet assay of cell lines transduced with sh scr or sh α#2. **(F)** Quantification of crystal violet stained area. **(G)** Immunoblot assessing PI5P4Kα and cleaved PARP protein levels in the indicated cell lines transduced with sh scr, sh α#1 or sh α#2 in the absence or presence of FAC. Tubulin was used as a loading control. Data is representative of three independent experiments. **(B, D, F** Data are presented as the mean ± sd. Statistical significance was calculated using unpaired two-tailed Student’s t-test without **(D,F)** or with Welch’s correc-tion **(B)**. ns – not significant, *P < 0.05, **P ≤ 0.001.

We then aimed to examine the underlying cause of the reduced intracellular iron levels. Intracellular iron production can occur via ferritinophagy, a form of selective autophagy^36,37^. Although we and others have established a role for PI5P4Kα in regulating autophagy flux in other biological contexts^14,38^, our analyses in PDAC cells indicated a decrease in levels of LC3-GFP, which gets restored in the presence of iron supplementation (Figures 5A, 5B and S3A). A decrease in LC3 levels alone is not a reliable indicator of whether autophagic activity is inhibited or stimulated; therefore, we probed autophagic flux using a mCherry-GFP-LC3 ratiometric assay^39^. We first validated this approach by employing Bafilomycin as a control, a known inhibitor of ferritinophagy^40^ and as expected Bafilomycin-induced loss of cell fitness is rescued by FAC supplementation (Figure S3B). Also, Bafilomycin increased levels of LC3-GFP (Figure S3C) and reduced the mCherry:GFP ratio indicative of reduced autophagic flux due to a block in the fusion of autophagosomes with lysosomes (Figures S3D and S3E). Next, we evaluated the mCherry:GFP ratio in the PI5P4Kα-depleted cells and we observed increased flux relative to control cells (Figures 5C-5F, S3F and S3G). This increase in autophagic flux was reversed by exogenous iron supplementation (Figures 5C-5F, S3F and S3G). Consistent with enhanced autophagy flux, we detected decreased levels of p62, an autophagic flux marker in PI5P4Kα knockdown cells (Figure 5G). These results indicate that the reduction in iron levels observed with PI5P4Kα knockdown was not due to autophagy inhibition^14^. Instead, autophagy was upregulated, likely as an adaptive response to intracellular iron deficiency. We then investigated the possibility that PI5P4Kα suppression could be affecting the import of iron into the cells. Recent studies have revealed interactions between the type I and type II PIPK enzymes, which have implications for membrane PI(4,5)P2 dynamics^41,42^. These interactions influence various cellular processes, including transferrin uptake, which is one of the major uptake mechanisms for iron acquisition. To investigate whether PI5P4Kα regulates transferrin uptake, which occurs through receptor-mediated endocytosis, we assessed transferrin internalization into the cells. Importantly, we observed a reduction in transferrin uptake upon depletion of PI5P4Kα (Figures 5H and 5I). These data show that PI5P4Kα depletion reduces intracellular labile iron pools by disrupting iron import into the cells and this imbalance in iron levels drives the apoptotic cell death.

**Figure 5.**
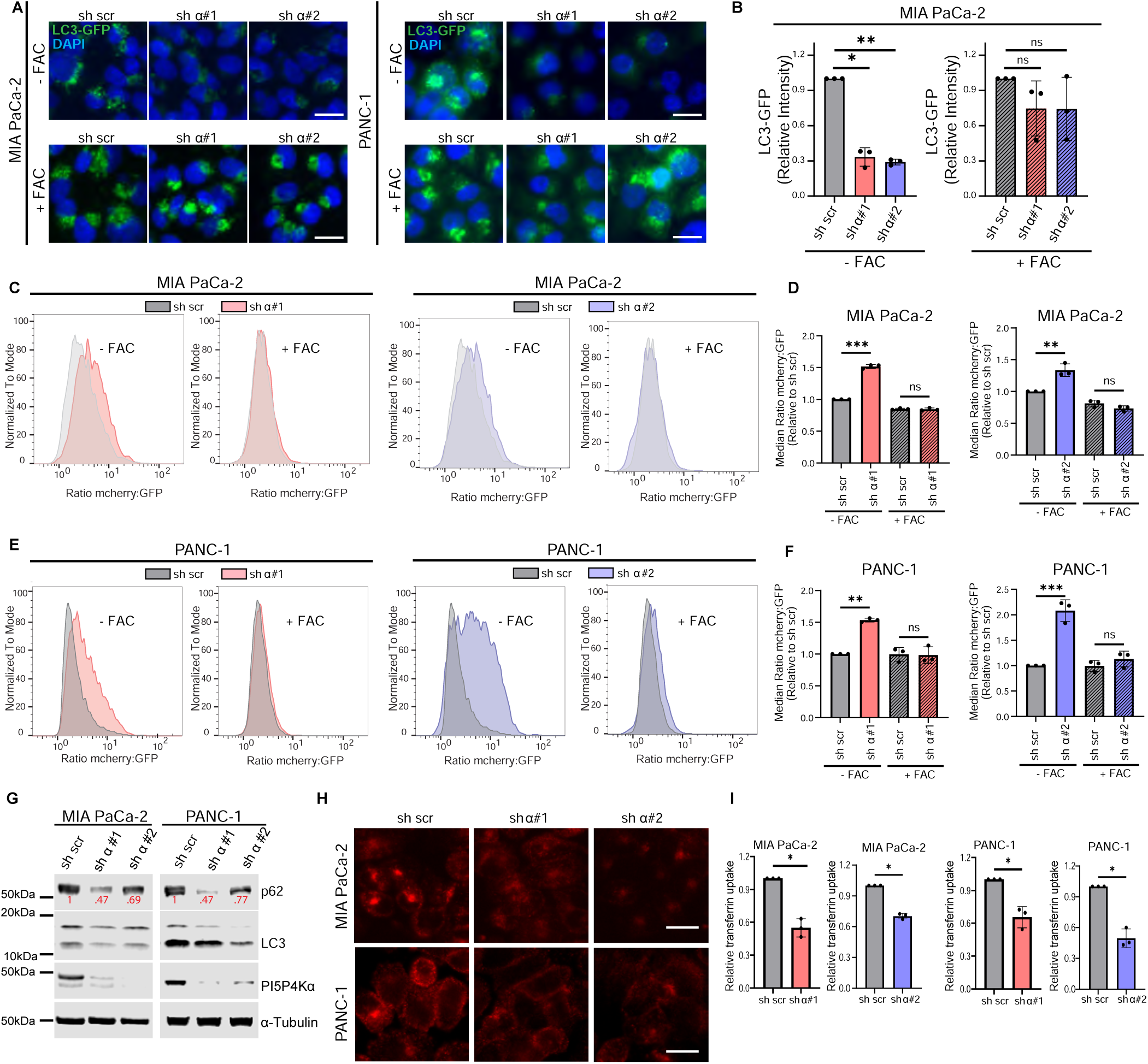
PI5P4Kα maintains intracellular iron levels via receptor-mediated endocytosis. **(A-B)** PDAC cells with stable expression of GFP-LC3 were transduced with sh scr, sh α#1 or sh α#2 in the absence or presence of FAC supplementation. Cells were imaged between 3-4 days post-transduction. **(A)** Immunofluores-cence microscopy images of LC3-GFP and DAPI indicating the representative effect of three independent experiments (scale bar, 20 µm). **(B)** Quantitation of immunofluorescence microscopy images in MIA PaCa-2 cells. Data is presented as signal intensity relative to the respective sh scr condition and is the average of three independent experiments. **(C-F)** PDAC cells with stable expression of mCherry-GFP-LC3 were transduced with sh scr, sh α#1 or sh α#2 in the absence or presence of FAC supplementation. Histogram plots generated from flow cytometry experiments indicating the ratio of mCherry to GFP as an assessment of the shift in autophagy flux. Histogram plots are representative of three independent experiments. Quantitation of the ratio of mCherry to GFP relative to sh scr. Data is the average of three independent experiments. **(C)** Histogram plots for MIA PaCA-2 cells. **(D)** Quantitation of the ratio of mCherry to GFP in MIA PaCa-2 cells. **(E)** Histogram plots for PANC-1 cells. **(F)** Quantitation of the ratio of mCherry to GFP in PANC-1 cells.**(G)** Immunoblot assessing PI5P4Kα, p62 and LC3 protein levels in the indicated cell lines trans-duced with sh scr, sh α#1 or sh α#2. Tubulin was used as a loading control. Red numbers indicate %knockdown over sh scr control after each condition is normalized to its respective Tubulin. Data is representative of three independent experiments. **(H-I)** Transferrin uptake in the indicated cell lines transduced with sh scr, sh α#1 or sh α#2. Cells were imaged between 3-4 days post-transduction. **(H)** Immunofluorescence microscopy images of Transferrin and DAPI indicating the representative effect of three independent experiments (scale bar, 20 µm). **(I)** Quantitation of immunoflu-orescence microscopy images. Data is presented as signal intensity relative to the sh scr condition and is the aver-age of three independent experiments. **(B, D, F, I)** Data are presented as the mean ± sd. Statistical significance was calculated using unpaired two-tailed Student’s t-test with Welch’s correction **(B, I)** and using one-way Anova **(D, F)**. ns – not significant, *P < 0.05, **P ≤ 0.001, ***P ≤ 0.0001.

### PI5P4Kα is overexpressed in human PDAC and promotes tumor growth in a mouse model

To examine the role of PI5P4Kα in human PDAC, we investigated the expression levels of *PIP4K2A* in human PDAC relative to normal pancreas tissue. We queried publicly available datasets including the TCGA dataset^43^, the GTEx portal and the NCBI Gene Expression Omnibus (GEO) database (GSE28735, GSE15471, and GSE42952). We observed a consistent upregulation of *PIP4K2A* transcript expression in the tumor samples relative to normal non-neoplastic tissue (Figure 6A) with overall survival showing poor outcome in patients with high expression of the *PIP4K2A* transcript^44^ (Figure 6B). To assess the expression of PI5P4Kα at the protein level we immunostained a PDAC tissue microarray, comprising samples from 121 patients who had undergone surgical resection. We found that PI5P4Kα protein was expressed in all 121 PDAC tumors and that 30/41 PDAC specimens demonstrated moderate or high levels of expression (Figure 6C). We then utilized the transcriptomics data from the TCGA portal and other datasets to investigate potential correlations between *PIP4K2A* expression and gene signatures associated with iron import. Intriguingly, we found that *PIP4K2A* expression levels positively correlated with the expression levels of genes associated with iron uptake and transport into the cells across multiple datasets (Figure 6D) and across different gene signatures (Figure S4A). Additionally, higher expression of the genes in the different iron import and transport signatures correlates with poor prognosis for PDAC patients (Figure 6E and S4B). These findings establish the expression profile of PI5P4Kα in human PDAC and support the concept that PI5P4Kα is linked to iron homeostasis.

**Figure 6.**
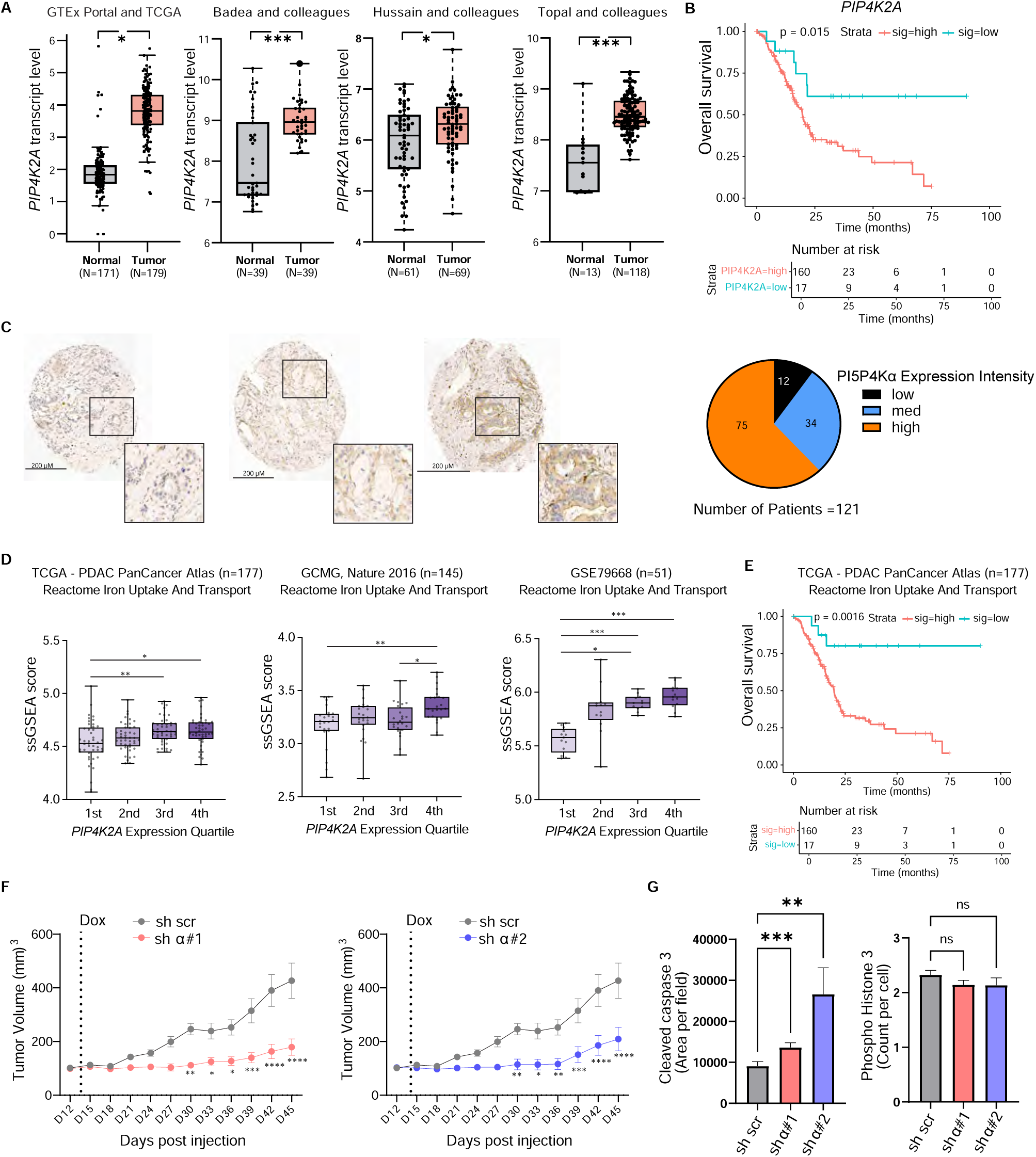
PI5P4Kα is highly expressed in human PDAC and promotes the growth of xenograft tumors. **(A)** *PIP4K2A* transcript levels in human PDAC tissues compared to normal pancreas or adjacent non-neoplastic tissue. Expression levels are depicted as box plots for the indicated datasets and include the information on the sample size (N). Statistical significance was calculated using unpaired two-tailed Student’s t-test. **(B)** Kaplan-Meier overall survival curve of PDAC patients stratified by *PIP4K2A* expression levels. Statistics was done using the Log-rank test. **(C)** Representative images of PI5P4Kα immunohistochemical staining of a tissue microarray (TMA) containing 121 surgically resected human PDAC samples. Patients were stratified into low, medium and high protein expression groups based on staining intensity and is represented on a pie chart (scale bar, 200µM). **(D)** Correlation plots between the expression of *PIP4K2A* from TCGA, GSE36924 and GSE79668 datasets with the Reactome gene signature of iron uptake and transport. **(E)** Kaplan-Meier overall survival curve of PDAC patients stratified by the expression levels of genes in the Reactome gene signature. **(F)** Heterotopic xenograft growth curves of MIA PaCa-2 cells expressing doxycycline (Dox)-inducible hairpins. The mice were put on Dox diet and were injected with doxycycline every two days, once the tumors reached approximately 100mm3 tumor volume (14 days post injection). Tumor volume measurements were done following Dox administration. Data is presented as the mean ± sem of n=7 (sh scr), n=6 (sh α#1) and n=9 (sh α#2). Statistical significance was calculated using two-way Anova. **(G)** Immunohistochemistry quantitation of tumor tissue sections with anti-cleaved caspase 3 and anti-phospho histone 3. For cleaved caspase 3, positive signal per image field was determined and for the pHis-H3, the number of p-HistoneH3-positive nuclei were normalized to the total cell count. Data is presented as the mean ± sem. **(D, G)** Statistical significance was calculated using non-parametric one-way ANOVA (Kruskal Wallis with Bonferroni correction). ns – not significant, *P < 0.05, **P ≤ 0.001, ***P ≤ 0.0001.

To investigate the role of PI5P4Kα in tumor growth control in the context of PDAC, we employed a doxycycline (Dox)-inducible shRNA system targeting *PIP4K2A* in MIA PaCa-2 cells. We verified that our Dox-inducible system leads to the reduction of PI5P4Kα expression (Figure S4C) and that this significantly suppresses PDAC cell fitness *in vitro* (Figure S4D). We used these cells to establish heterotopic xenograft tumors in athymic mice and 12 days post-inoculation when tumors reached an average volume of approximately 100 mm^3^, Dox administration was initiated (Figure S4E). Tumor measurements were obtained using a digital caliper every three days for 45 days post-implantation. We found that relative to the sh scr control tumors, knockdown of PI5P4Kα led to significantly reduced tumor growth (Figure 6F). Immunohistochemical analysis of the tumor sections revealed significantly elevated levels of apoptosis in the PI5P4Kα knockdown tumors, as assessed by staining with anti-cleaved caspase 3 (Figure 6G and S4F). To measure intratumoral proliferative capacity, we stained the tumors with anti-phospho histone H3, a marker of mitotic activity, which revealed that proliferation was unaffected by PI5P4Kα knockdown (Figure 6G and S4F). Collectively, these results demonstrate that PI5P4Kα plays a crucial role in supporting the growth of PDAC *in vivo*.

## Discussion

We identify PI5P4Kα as a critical metabolic dependency in PDAC and elucidate a novel role for PI5P4Kα in supporting their ferro-addiction^45,46^. While PI5P4Ks have been implicated in other cancers,^16^ their function in PDAC has not been previously reported. PI5P4Ks have been elucidated to support cancer metabolism but have never been shown to support iron homeostasis. PI5P4Ks do not usually exhibit mutational changes in cancers but their expression is often upregulated in tumor samples^16^, including in PDAC. This increase in expression potentially enables the PDAC cells to meet their increased nutrient demands, particularly in their nutrient-deprived metabolic environment, through the regulation of iron uptake and therefore intracellular iron levels. We found that the expression of *PIP4K2A* as well as of genes associated with iron uptake correlated with poor patient prognosis. Moreover, our analyses revealed a positive correlation between *PIP4K2A* expression and multiple gene signatures associated with iron uptake.

While investigating the mechanism of iron imbalance, we discovered that PI5P4Kα inhibition does not impair autophagy to limit the iron supply, as might have been expected based on previous studies in other models^14^ and from the recent studies on another PIP kinase family member, PIKfyve^47^ (cite). Instead, we observed an upregulation of autophagy following PI5P4Kα depletion, which was restored to basal levels with iron supplementation, indicating an adaptive response to reduced iron levels. The adaptive upregulation of autophagy has been a major concern since it drives resistance to therapies such as ERK and MEK inhibitors in PDAC^35,48^. With PI5P4Kα depletion, the increased autophagic activity does not defend against apoptosis. This highlights PI5P4Kα’s fundamental role in supporting overall metabolic homeostasis, emphasizing its potential as an effective therapeutic target. For regulating iron levels, we found that inhibition of PI5P4Kα results in a reduction in transferrin-bound iron uptake, potentially due to disruption in the phosphoinositide pools involved in regulating membrane dynamics, endocytosis, and vesicular trafficking^49–52^. PI5P4Ks primarily phosphorylate PI(5)P to generate PI(4,5)P2^9^ but has recently been shown to interact with the Type I PI4P5Ks to regulate plasma membrane PI(4,5)P2 levels^41^. Further, emerging research shows that PI5P4Ks can also utilize PI3P as a substrate to generate PI(3,4)P2^53^. Both these PIP2 species regulate endocytic trafficking pathways^50,54,55^ including the modulation of cellular iron uptake.

We show that depletion of PI5P4Kα significantly impairs the cellular fitness of PDAC cells, particularly those with TP53 mutations, leading to apoptotic cell death. This suggests that PI5P4Kα could serve as a therapeutic target in PDAC, especially in the context of p53 mutant status. Future investigations are required to elucidate the mechanism of this conditional essentiality of PI5P4Kα in p53-mutant PDAC cells. However, intracellular iron levels have been shown to be regulated by p53-dependent transcriptional and post-transcriptional regulation of genes involved in iron metabolism, providing a rationale for the sensitivity of p53-mutant PDAC cells to PI5P4Kα inhibition^56^. Importantly, the normal pancreatic epithelial cells were protected from PI5P4Kα-inhibition driven apoptosis, providing support for safe therapeutic utilization of PI5P4Kα inhibitors. The apoptotic cell death observed in PDAC cells following PI5P4Kα depletion is attributed to metabolic stress, stemming from the depletion of key TCA cycle intermediates, disrupted glutamine flux, and a diminished cytosolic iron pool. The ability to rescue cell fitness through the exogenous supply of TCA metabolites and iron further supports the idea that metabolic alterations are central to the vulnerability induced by PI5P4Kα inhibition in PDAC.

Overall, our study uncovers how PDAC cells leverage PI5P4Kα to support their iron addiction, positioning PI5P4Kα as a promising therapeutic target to improve outcomes for patients with this lethal cancer. With growing interest in developing PI5P4Kα inhibitors^1,16,18^, our findings suggest that transitioning from bench to clinic may soon be possible.

## STAR Methods

### Cells and cell culture conditions

MIA PaCa-2, PANC-1, SW1990 and HPAF-II were obtained from the American Type Culture Collection. MIA PaCa-2, PANC-1, SW1990 and HPAF-II were cultured in DMEM (10-017-CV, Corning), 10% FBS (89510-186, Avantor) and 20 mmol/L HEPES (11-330-057, Life Technologies). Media for hTERT-HPNE cells was supplemented with 10 ng/mL hEGF (E9644, Sigma). Cell lines were maintained under 5% CO2, 37°C in a humidified atmosphere in the presence of 100 U/mL penicillin/streptomycin (15140122, Life Technologies) and were routinely tested for mycoplasma contamination using the PCR Mycoplasma Detection Kit (ABM). For doxycycline-inducible expression system, the cell lines were selected and maintained in Tet System Approved FBS (631101, Takara). Lentiviral and retroviral transduced cells were selected in puromycin (NC9138068, Invivogen).

### Reagents

The following reagents were used: Polybrene (NC9840454, Santa Cruz Biotechnology), 37% Formaldehyde (RSOF0010, RICCA chemical company), Crystal Violet (C0775, Sigma), Emricasan (510230, MedKoo Biosciences), Tetramethylrhodamine Ethyl Ester Perchlorate (TMRE) (T669, Thermo Fisher Scientific), Human Transferrin, CF™594 Conjugate (00084, Biotium), DAPI (422801, BioLegend), Hoechst 33342 (0219030525, MP Biomedicals), Doxycycline hyclate (D9891, Sigma). For the supplementation assays with Sodium L-lactate (71718), Sodium Pyruvate (11360070, Thermo Scientific), Dimethyl α-ketoglutarate (DMKG, 349631, Sigma), Dimethyl DL-Glutamate hydrochloride (DMGlu, 18602-5, Cayman Chemical) Company), Ammonium iron (III) citrate (FAC) (F5879, Sigma), Ferrostatin (502259214, Fisher Scientific), sodium citrate (0101, VWR), the treatments were performed simultaneously with the transduction. For LC3-GFP imaging and autophagy flux experiments, cells were treated for 16hrs with 50nM of Bafilomycin A1 (S1413, Selleckchem). For doxycycline-inducible expression system, the transduced cells were treated for 3 days with 2 mg/mL of doxycycline for knock-down induction.

### Constructs, Viral Production and Transduction

The shRNA vector pLKO.1-puromycin was used to clone the human shRNA targeting PIP4K2A (sh α#1-TRCN0000220609; 5′-CCTCGGACAGACATGAACATT-3′ and sh α#2 - TRCN0000195145 ;5′-CCAGCATCGTTCTAGCTATTT-3′). They were cloned according to the Addgene shRNA cloning protocol. pLKO.1-puro sh scr was purchased from Addgene (17920). Tet-pLKO-puro constructs for doxycycline-induced shRNA expression were generated and used for cloning the human shRNA targeting *PIP4K2A*. They share the same targeting sequence as their pLKO-puro counterparts.

HEK293T packaging cell line was used for lentiviral and retroviral amplification. For lentiviral particle production, the lentiviral vectors were transfected along with the packaging plasmids psPAX2 (12260, Addgene,) and pVSVg, using Lipofectamine (11668019, Life Technologies). The packaging plasmid used for retroviral vector transfection was pCL-Ampho. The media was replaced the following day and virus-containing supernatant was collected after 48 hours by passing through a 0.45-μm filter, aliquoted, and stored at −80°C until further use. Cells were transduced with virus-containing media in the presence of 10μg/ml polybrene (Santa Cruz Biotechnology; NC9840454), prior to puromycin (NC9138068, Invivogen) selection.

### qRT-PCR

Total RNA was prepared using Directzol RNA MiniPrep (50–444-628, Zymo Research Corporation). cDNA was synthesized using High-capacity cDNA Reverse Transcription kit (4368814, ThermoFisher Scientific) and qRT-PCR performed utilizing PowerUp SYBR green (A25742, ThermoFisher Scientific) and the LightCycler 96 (Roche). Cells were plated in 6-well cell culture plates in complete media and transduced the next day with sh scr, sh α#1, and sh α#2 in complete medium. After 3 days, the RNA in the lysates was extracted using the Directzol RNA prep protocol. Relative target gene expression was determined by comparing average threshold cycles (CT) with that of housekeeping genes (*TBP* or *RPL13*) by the delta-delta CT method. The following primers were used in this study: PIP4K2A-F: CGTAGCGCAGAAAGTGAAGC and PIP4K2A-R: GGCTCGGCATGTTTTCTTTGT.

### Growth Curves and cell proliferation assay

The human PDAC cells and normal hTERT-HPNE cells were seeded in complete media on 96-well or 24-well cell culture plates. 24 hours after cell plating, a Day 0 plate was fixed to determine the relative cell number and the plates for the different time points were transduced with sh scr, sh α#1, and sh α#2. At the indicated time points, cells were fixed were fixed with 4% formaldehyde following which the wells were washed by submersion in a water bath. For crystal violet staining 0.5% crystal violet (C0775, Sigma) was added to the wells for 20 minutes. The solution was then removed, and the wells were thoroughly washed by submersion in a water bath. Plates were left to dry at 37°C and scanned and cell number was calculated as relative stained area determined using ImageJ.

For rescue assays with exogenous supplementation, the cells were transduced with sh scr, sh α#1, and sh α#2 in the absence or presence of Emricasan (30uM), pyruvate (1mM), lactate, dimethyl a-ketoglutarate (DMKG) (7 mM), dimethyl DL-glutamate (DMGlu) (15 mM), FAC (0.1mg/ml), sodium citrate (0.1mg/ml) and ferrostatin (1µM). The treated cells were fixed for crystal violet assay and cell number was calculated as relative stained area determined using ImageJ.

### Hoechst staining

Cells were seeded in 96 well plates in complete medium. 24 hours after seeding the cells, the cells were transduced with sh scr, sh α#1, and sh α#2 in complete medium. The cells were fixed with 4% formaldehyde and stained with 0.5 µg/ml Hoechst 33342 and 0.1% Triton X-100. The stained cells were imaged at 20× magnification using the EVOS FL Cell Imaging System (Thermo Fisher Scientific). Cells undergoing apoptosis demonstrate nuclear condensation (bright staining intensity) and were quantified in different fields of triplicate wells of three independent experiments and calculated as a percentage of total cell number.

### Immunoblot analysis

Total cell lysates were prepared by washing cells with ice-cold PBS. The cells were then lysed with RIPA lysis buffer, as well as Halt protease and phosphatase inhibitor cocktail (78440, Thermo Fisher Scientific,). Protein was measured using the BCA assay (23225, Thermo Scientific) and 20–30 μg of total cell lysates were run on an SDS–polyacrylamide gel electrophoresis. The proteins were transferred on to a nitrocellulose membrane and membranes were probed overnight at 4°C with the appropriate primary antibody. Antibodies used were as follows: PIP4K2α (5527, Cell Signaling;), PARP (46D11, Cell Signaling), β-actin (sc-517582, Santa Cruz Biotechnology;) and α-tubulin (T6199, Sigma), p62 (H00008878, Abnova), LC3 (12741, Cell Signaling). Membranes were incubated with appropriate secondary antibodies and visualized using a LI-COR Odyssey Imager.

### Autophagy Assay

Stable cell lines expressing the GFP-LC3 were plated on black flat bottom plate 96 well plate (6055302, Perkin Elmer Health Sciences) and transduced with sh scr, sh α#1, and sh α#2. The cells were serum starved overnight before being fixed with formaldehyde for 15 minutes. The wells were washed with PBS and the nuclei were stained with 2 mg/ml of DAPI(MilliPore). Images were captured at 40X magnification using the BioTek Cytation 5 (Agilent Technologies) from the Sanford Burnham Prebys Cell Imaging Core and were quantified using automated analysis using the BioTek Gen5 software. The signal intensity was normalized to the cell count.

Autophagy flux was measured using stable cell lines expressing the fusion protein mCherry-eGFP-LC3B, which distinguished autophagosomes (mCherry and GFP double positive) and autolysosomes (mCherry only). The stable lines were plated in 6 well plates in triplicate and the transduced cells were harvested and suspended in PBS. For each sample, 10,000 events were acquired on a SORP LSRFortessa (BD Biosciences, San Jose, CA) with FACSDiVa Software v8.0.2. Daily instrument QC was performed by SBP Core staff according to the manufacturer’s protocol. GFP was measured with 50 mW of 488 nm excitation and a 530/28BP emission filter; mCherry was measured with 50 mW of 561 nm excitation and a 610/20BP emission filter. Compensation was calculated by FACSDiVa using single-color GFP and mCherry controls along with a non-fluorescent parental cell line as the unstained control. Analysis was performed with FlowJo v10.10.0. Debris and aggregates were gated out, followed by an mCherry-high gate selecting cells more than 1 log brighter than the unstained control to avoid false positives due to increased autofluorescence from some of the experimental treatments. A derived parameter defined as compensated mCherry divided by compensated GFP was created. The gating images are included for all the experiments.

### Metabolite extraction and measurement

Cells were seeded in 6 well plates and were run in 3–5 replicate wells, and extra wells were used for cell counting and protein quantification for normalization. Three days after transduction, the cells were washed with PBS three times and then recovered by scraping into 450 uL of ice-cold methanol/ water (1:1 v/v), containing 20 µM L-norvaline as internal standard. Further extraction was performed by addition of 0.225 ml chloroform, with vortexing and centrifugation at 15,000×g for 5 min at 4 °C. The upper aqueous phase was dried under vacuum using a Speedvac, derivatized and analyzed using gas chromatography–mass spectrometry (GC–MS) to quantify small polar metabolites as described^57^. Fractional labeling from 13C substrates were calculated as described^57^. Data from standards was used to construct standard curves in MetaQuant^58^, from which metabolite amounts in samples were calculated. Metabolite amounts were corrected for recovery of the internal standard and for 13C labeling to yield total (labeled and unlabeled) quantities in nmol per sample, and then adjusted per cell number.

### Quantification of labile iron pool

The labile iron pool in the transduced cells was quantified using FerroOrange dye (F374-12, Dojindo,) that is highly selective for intracellular Fe^2+^ detection. Cells were plated in black flat bottom 96 well plate (6055302, Perkin Elmer Health Sciences). The transduced cells were incubated with 1 μM FerroOrange dye in serum free Fluorobrite (A1896701, Thermo Scientific) media supplemented with 4mM glutamine for 30 minutes. The wells were washed with the flurobrite media and imaged in that media. Images were captured at 40X magnification using the BioTek Cytation 5 (Agilent Technologies) from the Sanford Burnham Prebys Cell Imaging Core and were quantified using automated analysis using the BioTek Gen5 software. The signal intensity was normalized to the cell count.

### Transferrin uptake

Cells were plated in black flat bottom 96 well plate (6055302, Perkin Elmer Health Sciences). On the day of experiment the transduced cells were starved for 30 minutes in media without FBS at 37°C. Starved cells were prechilled on ice for 5 minutes and treated with prechilled Transferrin, CF™594 Conjugate (00084, Biotium) resuspended in serum-free media for 30 minutes on ice. The media was then replaced with prewarmed media containing FBS, and cells were incubated at 37°C for 10 minutes to initiate transferrin uptake. Cells were then washed twice with PBS, fixed using 4% PFA for 15 minutes and stained with DAPI^35^. Images were captured at 40X magnification using the BioTek Cytation 5 (Agilent Technologies) from the Sanford Burnham Prebys Cell Imaging Core and were quantified using automated analysis using the BioTek Gen5 software. The signal intensity was normalized to the cell count.

### Analysis of human PDAC datasets

*PIP4K2A* gene expression data from human PDAC public datasets were used for tumor versus normal comparison. Data were obtained from The Cancer Genome Atlas, GTEx portal and the Gene Expression Omnibus (accession numbers GSE28735, GSE15471, GSE42952). Correlation plots were generated between the expression of *PIP4K2A* with either the GSEA iron import into the cell gene signature or the Reactome gene signature of iron uptake and transport. For the correlation plots data was used from TCGA, GSE36924 and GSE79668.

### Animal studies

All animals were housed in sterile caging and maintained under pathogen-free conditions. All animal handling and experimental procedures were performed under the Institutional Animal Care and Use Committee–approved protocol. Relative humidity in the rodent rooms was monitored and recorded daily by animal technicians but was not controlled. Generally, the relative humidity of the animal room is within the acceptable range of 30–70% for rodents. A standard diurnal light cycle of 12 h light, 12 h dark was used, with a rodent room thermostat temperature set point of 72 °F. 1 × 10^6^ stably transduced MIA PaCa-2 cells, were suspended in 100 µl of 1:1 PBS/Matrigel (354234, Corning) and injected subcutaneously into the flanks of female nude mice (Homozygous Fox1nu, Jackson Laboratory, 8 weeks old). Palpable tumors were used for digital caliper measurements in two dimensions to estimate tumor volume according to the equation *V* = (*L* × *W*^2^)/2. For doxycycline administration, mice were treated when tumors reached approximately 100 mm^3^. Doxycycline was administered through IP injection every 2 days and through diet (TD.01306, Envigo). Tumor sizes were recorded by electronic caliper measurement and tumors were collected and cut into pieces when reaching a volume of approximately 1,000 mm^3^. For immunoblots, the samples were snap-frozen in liquid nitrogen.

For IHC, the cross-section tumor slices were fixed in 10% formalin and fixed tissue was embedded in paraffin and sectioned by the histology core at SBP. Detection was done using the VECTASTAIN Elite ABC HRP Kit (Vector Labs) and the DAB HRP Substrate Kit (Vector Labs). For nuclear counterstaining, sections were stained with hematoxylin. Following dehydration, the slides were mounted with coverslips and Permount mounting medium (Fisher Scientific). Images were captured with a brightfield Olympus CX-31 microscope coupled with INFINITY camera and INFINITY capture software (Lumenera). The following primary antibodies and dilutions were used: p-Histone H3 (1:200) (9701, Cell Signaling) and cleaved caspase 3 (1:1000) (9664, Cell Signaling) For the pHis-H3, the number of p-HistoneH3-positive nuclei normalized to the total cell count and for cleaved caspase 3, positive signal per image field was determined using the Fiji software (NIH).

### TMA and IHC detection

All work including human specimens was conducted under the ethical approval (BASEC-2021-00417), all patients further signed a general consent to agree to use their tissue for research. A 2μm thick tissue section from the self-made PDAC tissue micro array was deparaffinized in xylene, rehydrated using decreasing concentrations of ethanol (100 to 95 to 70% to H20). Heat-induced epitope retrieval was done at 95°C for 30 min in Tris-EDTA buffer. The entire staining procedure was conducted with the Ventana Benchmark semi-automated staining system using Ventana reagents for the entire procedure (including iVIEW DAB detection kit). Incubation with primary antibody PIP4K2A (Proteintech, 1249–1-AP) was done at a 1:100 dilution.

### Quantification and statistical analysis

All graphs were made using GraphPad Prism software (GraphPad). Results are shown as the mean of at least three independent experiments ± standard deviation (SD) unless stated otherwise. Statistical significance was determined by the unpaired two-tailed Student’s t test with Welch’s correction when appropriate. One-way Anova was used for analysis when all groups are compared to sh scr and for correlation analysis of *PIP4K2A* expression with iron gene signatures. Two-way Anova was used for tumor growth analysis. P values less than 0.05 were considered statistically significant (*p < 0.05, **p ≤ 0.01, ***p ≤ 0.001 and ns=not significant).

## Acknowledgements

We extend our gratitude to the following cores at SBPMDI (NCI P30 CA 030199): Cell Imaging, Histology, Flow Cytometry and Bioinformatics. We also appreciate the valuable feedback and discussions from the Emerling and Commisso laboratories. Special thanks to the Sanford Burnham Prebys Cancer Metabolism Core for conducting the 13C-labeling and polar metabolite quantification experiments, and to Guillermina Garcia and Monica Sevilla at the Sanford Burnham Prebys Histology Core for preparing the samples for histology. The graphical abstract was created using BioRender.com. D.A.S was supported by NCI Research Specialist Award R50 CA283813. This work was funded by NCI (R01 CA 237536), NIGMS (R01 GM143583), and ACS (RSG-20-064-01-TBE) to B.M.E. This work was also supported by NIH grants R01 CA254806 and R01 CA207189 to C.C. Sanford Burnham Prebys Medical Discovery Institute core services are supported by NCI Cancer Center Support grant P30CA030199.

## Contributions

G.K.A., B.M.E and C.C. designed the study, wrote the manuscript and prepared all the figures. G.K.A., R.L., K.L., C.M.G., A.L. and T.A., performed experiments and/or analyzed data. C.P performed the TMA staining. D.A.S. ran the metabolomic samples. Y.A analyzed flow cytometry experiments. R.M performed the bioinformatic analysis. B.M.E and C.C. supervised the study. All authors reviewed, edited or commented on the manuscript.

## Supplemental Information

Document S1. Figures S1-S4.

---

**Figure S1.**
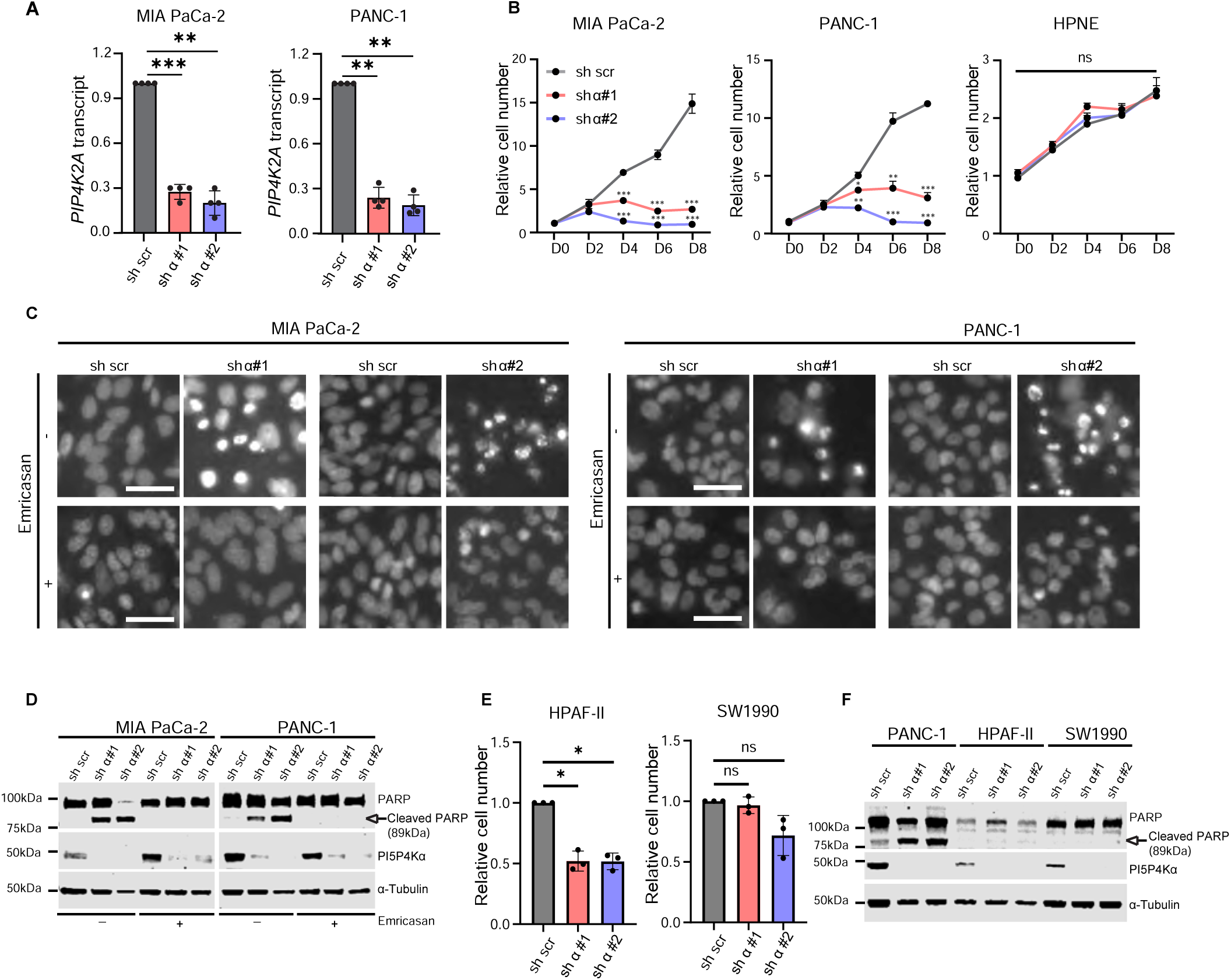
PI5P4Kα depletion induces apoptosis selectively in p53-mutant PDAC cells. **(A)** Relative *PIP4K2A* mRNA levels as assessed by qPCR in MIA PaCa-1 or PANC-1, 3 days post-transduction with sh scr, sh α#1 or sh α#2. Data are presented relative to sh scr and are average of four independent experiments. **(B)** Quantification of relative cell number using the crystal violet stained area for each cell line at the indicated days post-transduction. Data is presented relative to the sh scr condition and is representative of three independent experiments. **(C-D)** The indicated cell lines were transduced with sh scr, sh α#1 or sh α#2 in the absence or presence of Emricasan. **(C)** Representative images of Hoechst staining. (scale bar, 50µM)**. (D)** Immunoblot assessing PI5P4Kα and cleaved PARP protein levels in the indicated cell lines. Tubulin was used as a loading control. Data is representative of three independent experiments. **(E)** Quantification of crystal violet stained area in HPAF-II and SW1990 cell lines on D8 post-transduction. Data is presented relative to the sh scr condition and is the average of three independent experiments. **(F)** Immunoblot assessing PI5P4Kα and cleaved PARP protein levels in the indicated cell lines. Tubulin was used as a loading control. Data is representative of three independent experi-ments. **(A, B, E)** Data are presented as the mean ± sd. Statistical significance was calculated using unpaired two-tailed Student’s t-test with Welch’s correction. ns – not significant, *P < 0.05, **P ≤ 0.001, ***P ≤ 0.0001.

**Figure S2.**
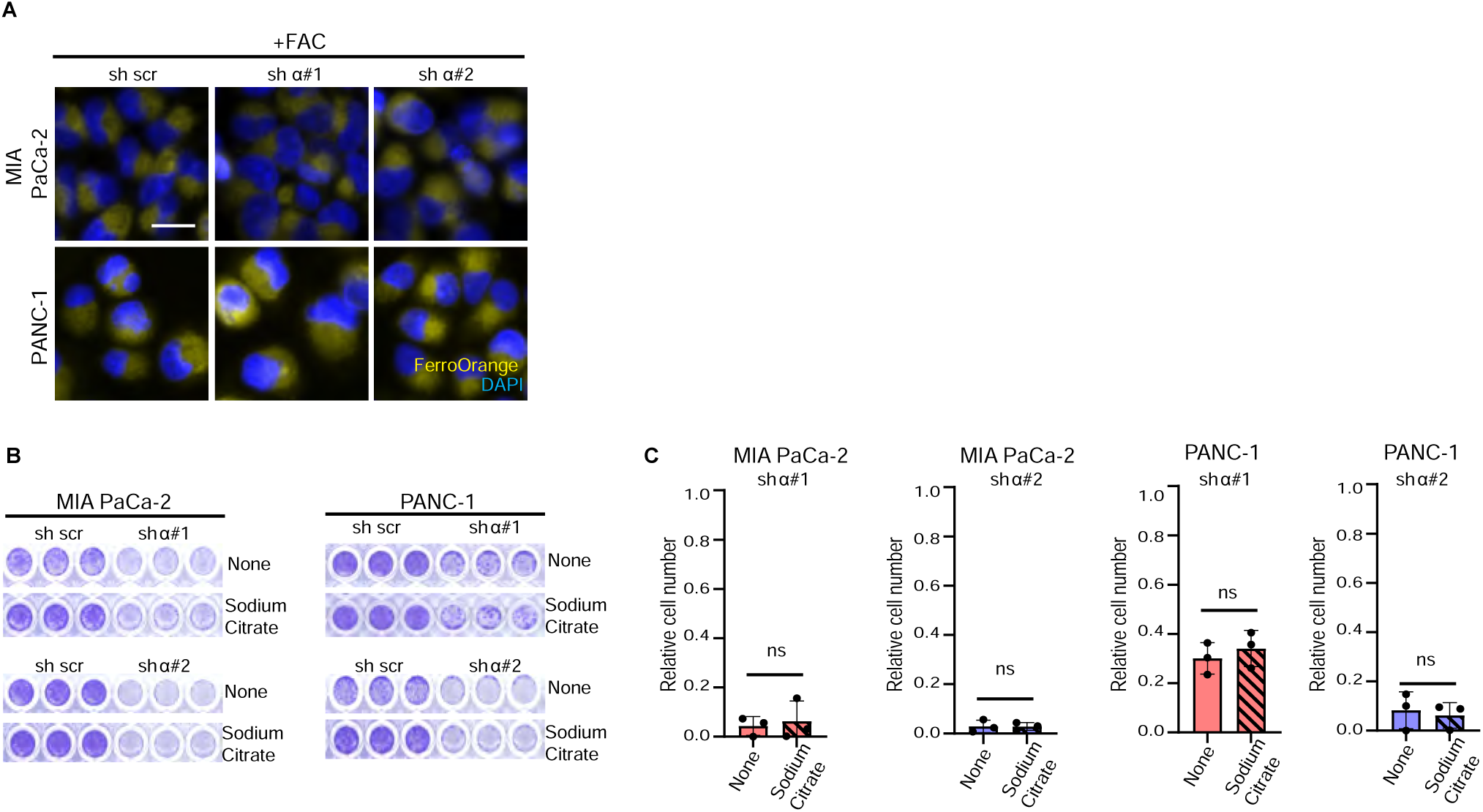
PI5P4Kα is a critical regulator of cellular iron levels. **(A)** Immunofluorescence microscopy images of FerroOrange staining in the indicated cell lines transduced with sh scr, sh α#1 or sh α#2 in the presence of FAC supplementation. Cells were imaged between 3-4 days post-transduction. Data is the representative effect of three independent experiments (scale bar, 20 µm). **(B-C)** Crystal violet staining and quantification of relative cell number in the indicated cell lines transduced with sh scr, sh α#1 or sh α#2 in the absence or presence of sodium citrate. **(B)** Representative image of crystal violet assay. **c,** Quantification of crystal violet stained area. Data are presented as the mean ± sd. Statistical significance was calculated using unpaired two-tailed Student’s t-test. ns-not significant.

**Figure S3.**
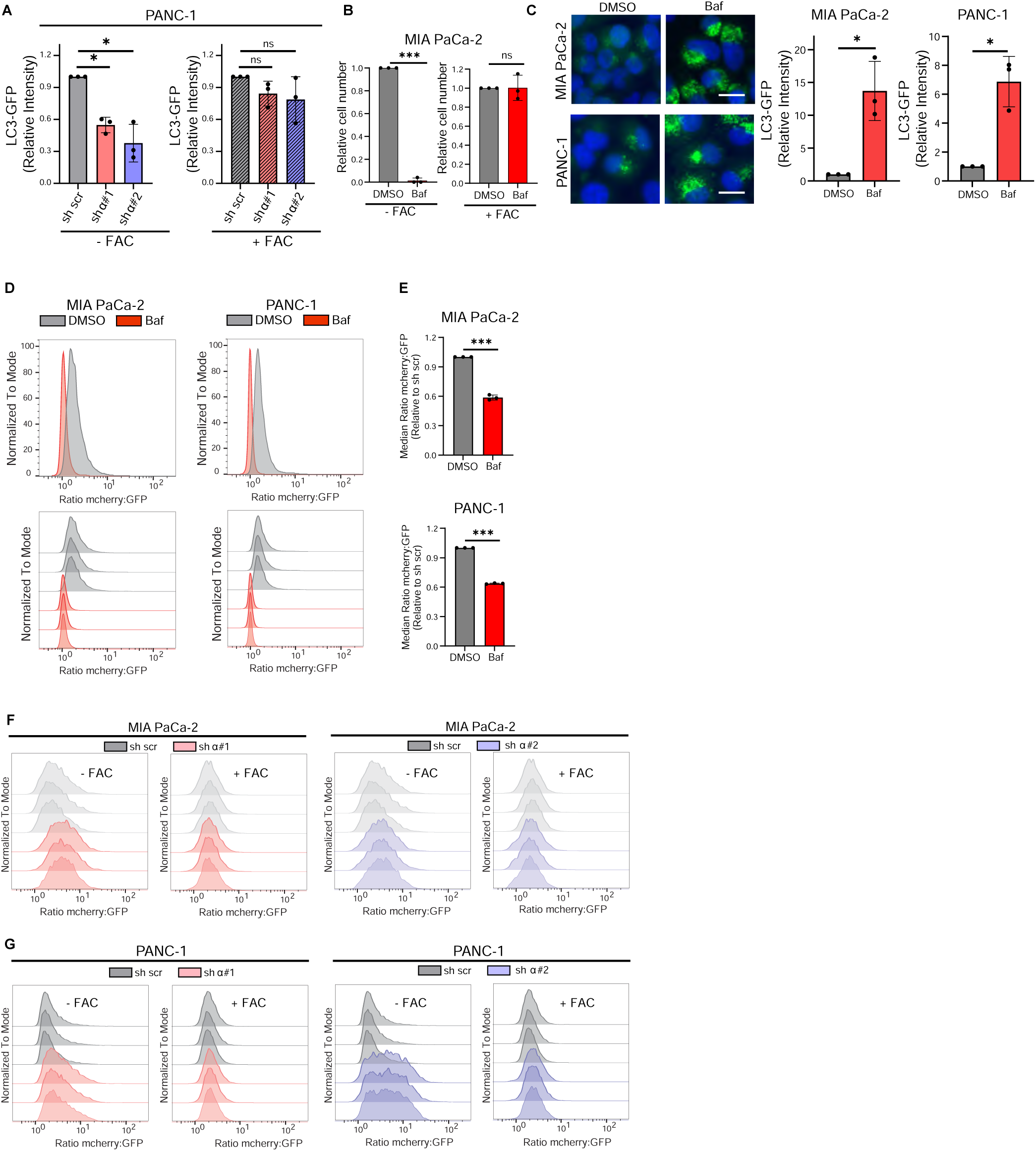
PI5P4Kα depletion triggers an adaptive autophagic response. **(A)** PDAC cells with stable expression of GFP-LC3 were transduced with sh scr, sh α#1 or sh α#2 in the absence or presence of FAC supplementation. Cells were imaged between 3-4 days post-transduction. Quantitation of immuno-fluorescence microscopy images in PANC-1 cells. Data is presented as signal intensity relative to the respective sh scr condition and is the average of three independent experiments. **(B)** Quantification of crystal violet stained area in cells treated with DMSO and Bafilomycin (Baf) 5nM in the absence or presence of FAC supplementation.**c,** Immuno-fluorescence microscopy images of LC3-GFP and DAPI in cells treated with DMSO and Baf (50nM), indicating the representative effect of three independent experiments (scale bar, 20 µm). Quantitation of immunofluorescence microscopy images. Data is the average of three independent experiments. **(D, E)** PDAC cells with stable expression of mCherry-GFP-LC3 were treated with DMSO or Baf (50nM) for 16hrs. **(D)** Histogram plots generated indicating the ratio of mCherry to GFP. Histogram plots are representative of three independent experiments. **(E)** Quantitation of the ratio of mCherry to GFP relative to sh scr. Data is the average of three independent experiments. **(F, G),** PDAC cells with stable expression of mCherry-GFP-LC3 were transduced with sh scr, sh α#1 or sh α#2 in the absence or pres-ence of FAC supplementation. Histogram plots of the flux assay showing the individual replicates of the representa-tive experiment in MIA PaCa-2 cells **(F)** and PANC-1 cells

**Figure S4.**
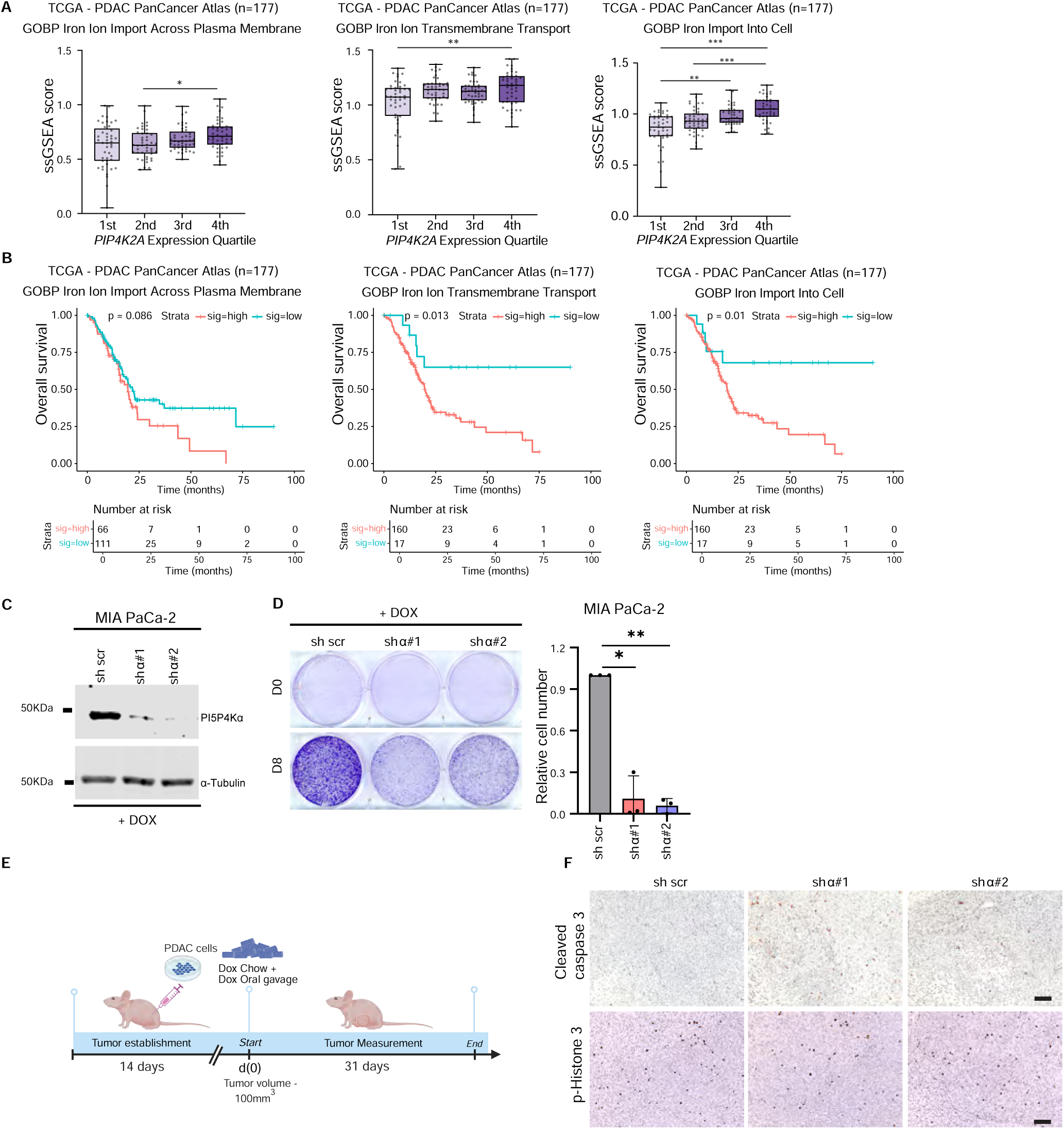
*PIP4K2A* gene expression correlates with iron import gene signatures. **(A)** Correlation plots between the expression of *PIP4K2A* from TCGA and GOBP iron import across plasma membrane and GOBP iron ion transmembrane transport and GOBP Iron import into cell gene signatures. **(B)** Kaplan-Meier overall survival curve of PDAC patients stratified by the expression levels of genes in the same gene signatures as in **(A)**. **(C, D)** MIA PaCa-2 cells with stable expression of doxycycline-inducible sh scr, sh α#1 or sh α#2 were treated with doxycycline. **(C)** Immunoblot assessing PI5P4Kα after treating the cells with doxycycline for 3 days. Tubulin was used as a loading control. Data is representative of three independent experiments. **(D)** Relative cell number as quantified by the intensity of the crystal violet stained area, eight days post doxycycline treatment. Data is normalized to respective D0 and is presented relative to the sh scr condition and is the average of three independent experiments. Data is presented as the mean ± sd. **(E)** Schematic representation of the in vivo xenograft study showing the timeline of tumor growth. **(F)** Immunohistochemistry images of tumor tissue sections with anti-cleaved caspase 3 and anti-phospho histone 3. Scale bar, 100 µm.Statistical significance was calculated using with non-parametric one-way ANOVA **(A)** and unpaired two-tailed Student’s t-test with Welch’s correction **(D)**. *P < 0.05, **P ≤ 0.001.

